# uBin – a manual refining tool for metagenomic bins designed for educational purposes

**DOI:** 10.1101/2020.07.15.204776

**Authors:** Till L.V. Bornemann, Sarah P. Esser, Tom L. Stach, Tim Burg, Alexander J. Probst

## Abstract

Resolving bacterial and archaeal genomes from metagenomes has revolutionized our understanding of Earth’s biomes, yet producing high quality genomes from assembled fragments has been an ever-standing problem. While automated binning software and their combination produce prokaryotic bins in high-throughput, their manual refinement has been slow and sometimes difficult. Here, we present uBin, a GUI-based, standalone bin refiner that runs on all major operating platforms and was specifically designed for educational purposes. When applied to the public CAMI dataset, refinement of bins was able to improve 78.9% of bins by decreasing their contamination. We also applied the bin refiner as a standalone binner to public metagenomes from the International Space Station and demonstrate the recovery of near-complete genomes, whose replication indices indicate active proliferation of microbes in Earth’s lower orbit. uBin is an easy to install software for bin refinement, binning of simple metagenomes and communication of metagenomic results to other scientists and in classrooms. The software is open source and available under https://github.com/ProbstLab/uBin.

## Main Text

Genome-resolved metagenomics aims at recovering genomes from shotgun sequencing data of environmental DNA. The genomes allow determination of the metabolic capacities of the individual community members and provide the basis for many downstream ‘omics techniques like metatranscriptomics and metaproteomics. Results from these technologies can provide important insight into the interactions of microbes within the community and with the environment [1,2]. While long-read sequencing can nowadays produce complete genomes from environmental samples [2], the percentage of closed genomes from complex ecosystems remains, however, as low as 5.3% [3]. Consequently, genomes need to be binned from metagenomes using genome-wide shared characteristics like their similar abundance pattern and k-mer frequencies [4,5]. Many automatic and semi-automatic tools have been developed to extract genomes from metagenomes [6–10]. The quality of the resulting bins, however, can vary greatly depending on metagenome complexity, sample type or microbial community characteristics [6]. Recent studies have shown that contamination in genomes from metagenomes in public databases is a frequent occurrence [11,12] and suggested genome curation as a mandatory analysis step prior to genome submission to public databases [13].

While established tools exist to determine the bin quality [6,14], i.e. searching candidate genomes for ubiquitous or specific marker genes to evaluate completeness and contamination, tools to improve upon the bin quality are sparse. Some established tools are used for genome refinement [15,16] but have not been designed for educational purposes and are sometimes not open source [16]. Consequently, we developed uBin as an interactive graphical-user interface that is easy to install on Mac OS, Windows, and Ubuntu for usage in, e.g., classrooms. uBin is inspired by ggKbase [16] and enables the curation of genomes based on a combination of GC content, coverage and taxonomy and couples this to information on completeness and contamination for supervised binning. In addition, uBin can be directly used as a standalone software to bin genomes from low complexity samples.

We tested the performance of uBin (MacOS, 16 GB of RAM) on simulated datasets with varying complexity of the Critical Assessment of Metagenome Interpretation (CAMI) challenge. The pre-assembled CAMI scaffolds were binned using four automated binners (using tetranucleotide frequency and differential coverage) and the results were aggregated using DAS Tool [6] (see Supplementary Methods for details). The dereplicated bins were curated using uBin, and the quality of the bins before and after curation was compared to the correct assignment based on the CAMI dataset (see *Tab. S1* for F-scores of Bins pre- and post-uBin curation). uBin curated bins showed a highly significant quality improvement in medium (p < 10^−4^) and high complexity datasets (p < 10^−5^), using both paired t-test and unpaired Kruskal-Wallis tests (*Fig. 1A*). No significant difference could be detected for the low complexity dataset (p > 0.70 / 0.65).

**Fig. 1.**
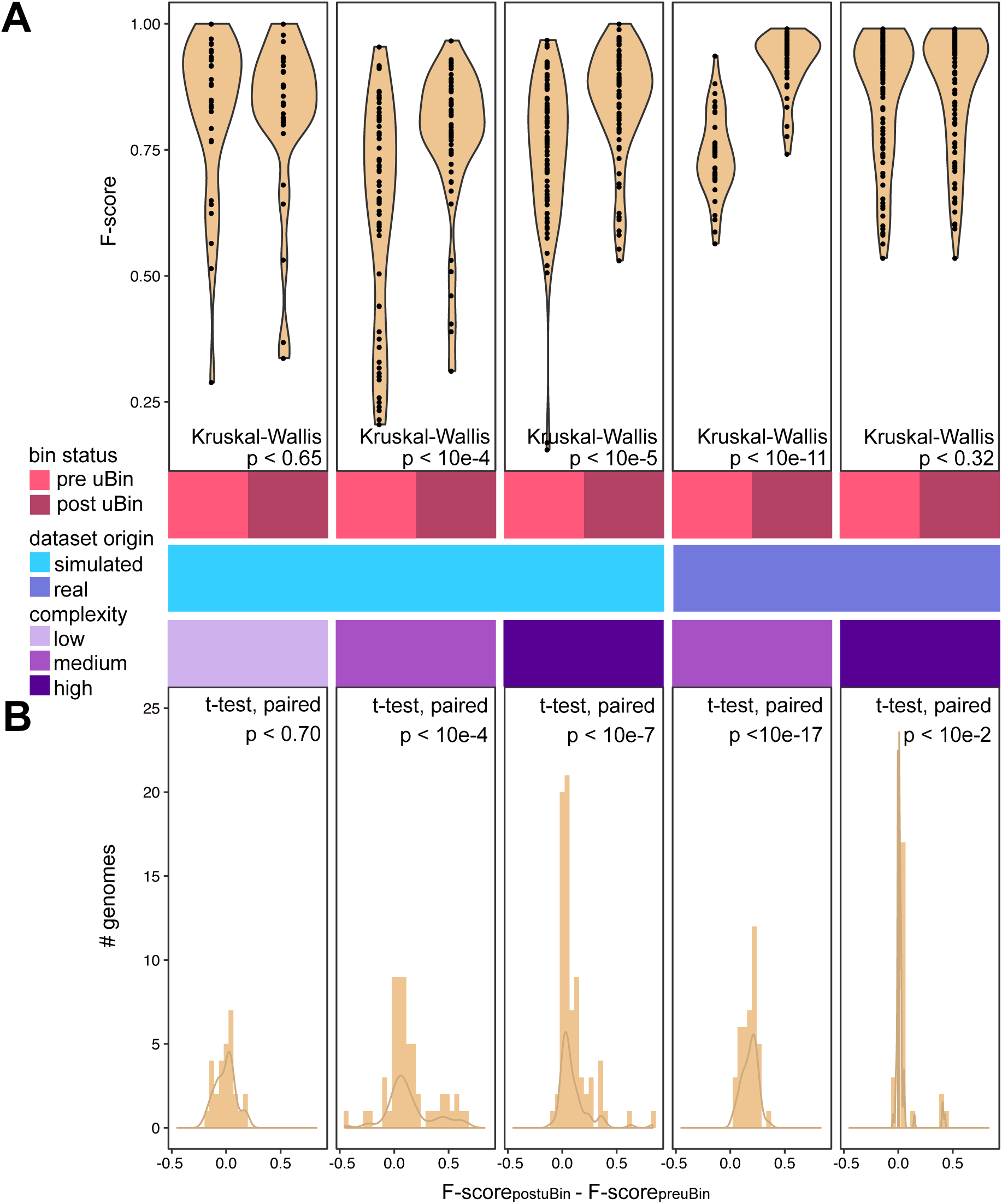
Performance of uBin on simulated and real datasets with varying degrees of complexity. **A:** Violin plots of the F-score (mean between recall and precision) of genomes prior to uBin curation (pre uBin) and after uBin curation (post uBin) across simulated low, medium and high complexity datasets of the CAMI challenge as well as real world metagenomic datasets of medium (Tomsk) and high (SulCav AS07-7) complexity. Unpaired Kruskal-Wallis p-values are depicted. **B:** Histograms of the F-score differences for each bin prior to and post uBin curation and their density distribution. Paired Welch t-test p-values are shown.

The bin quality of the low complexity dataset was significantly higher than the bin quality in medium (0.197 higher F-score, p < 10^−6^) and high complexity (0.118 higher F-score, p < 10^−4^) datasets (ANOVA coupled to TukeyHSD, p < 2×10^−6^) after DAS Tool [6] bin aggregation. Subsequent to curation with uBin the differences between these datasets were much less pronounced (ANOVA, p < 0.01), with only the high to medium complexity dataset showing a significant difference (p < 0.01, average 0.077 higher F-score in high complexity). We conclude that low complexity datasets bin very well with automated binners, while medium to high complexity datasets can greatly benefit from manual curation.

To challenge the above-mentioned conclusion, we applied uBin for the curation of bins from environmental metagenomes of medium and high complexity. As the true genome composition is unknown for these datasets, we used CheckM [14] to assess the completeness and contamination of constructed genomic bins. CheckM [14] is an independent metric compared to the marker sets used within DAS Tool [6] and uBin (see *Tab. S1* for F-scores of bins pre- and post-curation). We detected a significant improvement in genome quality when using uBin curation and directly comparing the bins in paired tests (p-values are provided in *Fig. 1B*).

Following the conclusion that binning of low complexity genomes can be achieved easily, we tested uBin’s capability as a standalone binner compared to Emergent-Self-Organizing Maps (ESOMs) [8] on public metagenomes of the International Space Station (ISS). uBin outperformed ESOM-based binning when used as a standalone tool and when used as a curation tool of the ESOM bins (*Fig. 2A*, see Supplementary Material for details). Using uBin, we successfully reconstructed 53 genomes with at least 94 percent completeness (*Fig. 2B*) and only 6% or less contamination (see *Tab. S2* for completeness and contamination statistics of recovered ISS genomes). When comparing their phylogenetic placement based on 16 ribosomal proteins to the taxonomic classification of uBin, we observed agreement between the taxonomic classification methods (see *Tab. S3* for the phylogenetic and uBin-based taxonomic placement of genomes). The one exception was the genome ISS_JPL_2332_S1_L003_Corynebacterium_afermentans_66_84, which was phylo-genetically placed next to a *Turicella* genome [17]. This genome has since been reclassified as *Corynebacterium otitidis* ATCC 51513 (NZ_AHAE00000000, see File S1 for the full phylogenetic tree).

**Fig. 2.**
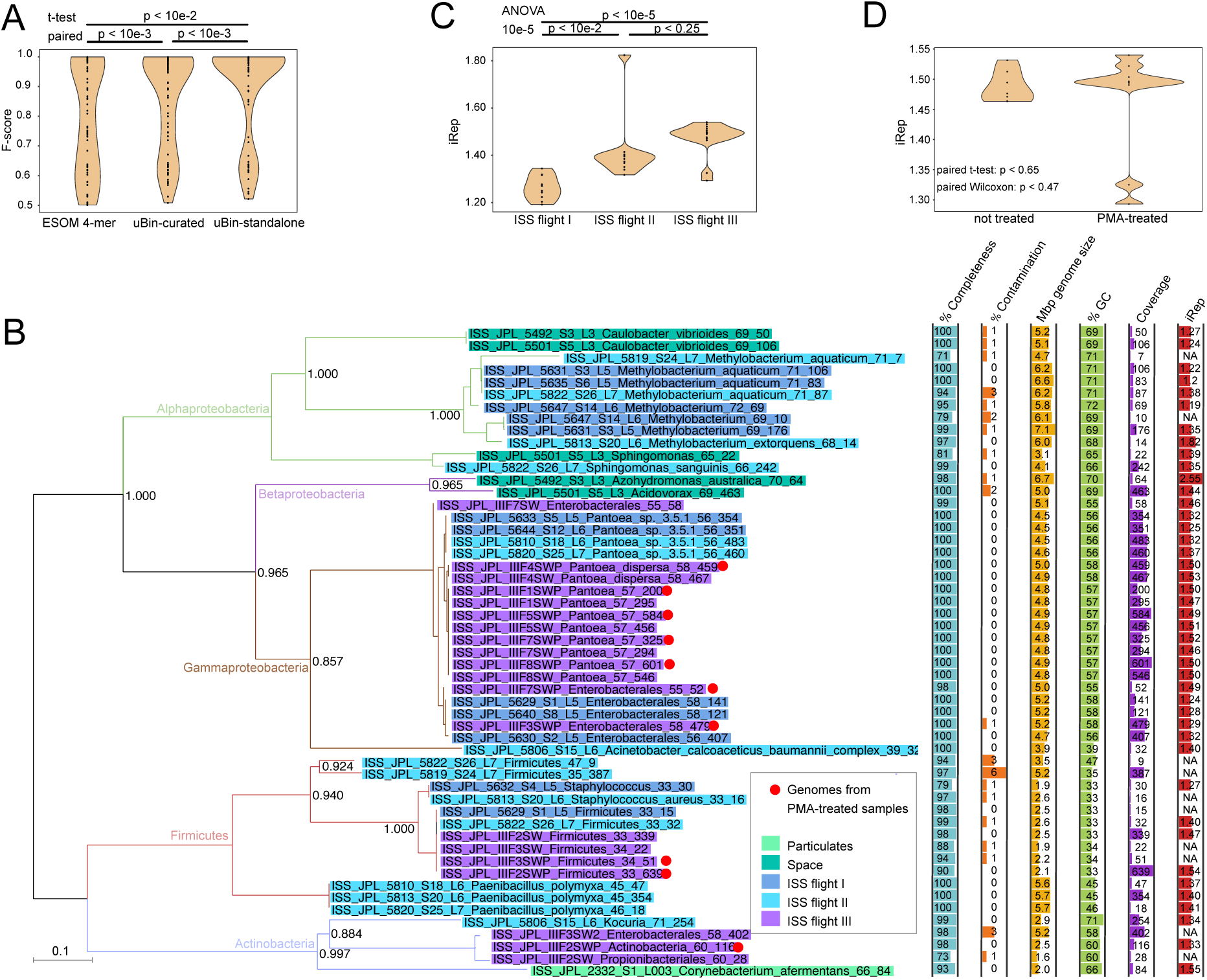
Reconstruction of genomes from the ISS, scoring of their curation and their phylogeny. **A:** Comparison of genome statistics after ESOM 4-mer binning, after uBin curation and after standalone binning using uBin. p-values correspond to paired Welch t-tests. **B:** Phylogenetic reconstruction based on the concatenation of 16 ribosomal proteins of 53 genomes from ISS metagenomes when using uBin as standalone binner. Branch colors indicate phyla assignments with coloring of leaves on tree displaying the sampling origin of the genomes. Genomes from PMA-treated samples (see main text) are highlighted with a red circle. The bargraphs on the right panel display completeness, contamination, genome size, GC content, coverage (relative abundance based on read-mapping) and the *in situ* replication measure (iRep [1]). **C:** Replication index dependency on flight of origin and significance testing thereof using ANOVA followed by TukeyHSD. **D:** Effect of PMA-treatment for removal of extracellular DNA on iRep of genomes from PMA-treated samples having increased iRep variance but no significant differences in iRep value based on paired Wilcoxon and paired t-tests (n=7 per group). Genomes were paired based on sample ID as well as using their shared uBin-taxonomy and GC content.

These bins represent an important step for space science since these are the first environmental genomes reconstructed from the ISS or associated transport flights. To investigate if the genomes are actively replicated under these conditions, we calculated the *in situ* replication measure iRep [1] for 43 out of 53 genomes. Across all sampling sites, the replication rates of the recovered population genomes varied from 1.20 to 2.55, which implies an active metabolism. For instance, the lowest iRep value, which was calculated for *Methylobacterium aquaticum*, indicated that on average 20% of its sampled population was undergoing genome replication. While closely related organisms often had similar replication measures (*Fig. S3*), the main discriminatory factor for varying replication indices was the origin of the flight (*Fig. 2C*) indicating community-wide shifts in replication between the different flights. The dataset also enabled the answer to a long-standing question of indoor microbiology relating to how external DNA influences the measurements of iRep values in metagenomics. Samples of the third sampled ISS flight were analyzed using both regular metagenomics as well as metagenomics following propidium monoazide (PMA) treatment, which removes external DNA fragments and enables DNA sequencing of cells with intact membranes. When comparing the iRep values of the paired samples (n=7 per group), no significant difference could be observed (paired t- and Wilcoxon-tests, *Fig. 2D*), although the variance of the iRep values increased tremendously after PMA treatment. Equivalence testing confirmed that there are no differences between these two sample types (p < 0.01). We suggest that PMA-treatment can improve the accuracy of iRep measures of environmental samples and recommend its usage where appropriate.

The herein presented uBin software is designed for improvement of bins and as a standalone binner for simple metagenomes with few species. It is independent of the operating system (available for Windows, MacOS, Linux) and GUI-based so that a wide audience of non-bioinformaticians can make use of it. The initial data processing (as general metagenomic data processing) necessitates bioinformatics knowledge but respective easy-to-use wrapper scripts are provided along with the software. Thus, uBin is ideally used by bioinformaticians to communicate metagenomic data to non-bioinformatics peers and to students in classrooms. After binning or curation with uBin, the user can deploy each genome into individual fasta files. These genomes can then be further explored for metabolic analyses with, e.g., MAGE [18] or KEGG mapper [19]. Consequently, uBin represents an important software link between automated binners along with the widely-used software DAS Tool and downstream analyses including genome refinement to completion [20].

## Acknowledgments

This study was funded by the Ministerium für Kultur und Wissenschaft des Landes Nordrhein-Westfalen (“Nachwuchsgruppe Dr. Alexander Probst”). We thank the students who tested and worked with uBin over the last two years in classrooms. We thank Christine Sun for her contribution to the script for calculating consensus taxonomy of scaffolds and Kasthuri Venkateswaran for input regarding sampling locations of the ISS samples. We thank Ken Dreger for the administration and maintenance of our servers.

## Supplementary Material for

### Supplementary Methods

#### Software implementation

uBin is written in TypeScript(3.2+)/JavaScript. It utilizes React (https://reactjs.org/) for its user interface and Redux (https://redux.js.org/) to manage the application state/data.

All imported data is stored in a local SQLite (sqlite3) database. Communication between uBin and the database is abstracted through TypeORM (https://typeorm.io/), an ORM written in TypeScript. To build the application and to provide cross-platform support, we use Electron (https://www.electronjs.org/).

The user interface uses HTML/CSS + Blueprint JS (a User-Interface (UI) toolkit, https://blueprintjs.com/) for general UI elements, react-vis (https://uber.github.io/react-vis/) for its Sunburst plot, and VX (a library for d3-based React visualization components, https://github.com/hshoff/vx)for every other plot. Crossfilter (https://github.com/crossfilter/crossfilter) is used to calculate the data to be plotted on-the-fly.

#### Metagenomic data assembly and processing

Quality control of ISS metagenome raw reads was performed using BBduk (B Bushnell, http://jgi.doe.gov/data-and-tools/bb-tools/) and Sickle [21]. Reads were assembled into contigs and scaffolded using metaSPAdes 3.12 [22] (see *Tab. S5* for read and assembly statistics). Genes were predicted for scaffolds larger than 1 kbp using Prodigal [23] in meta mode and annotated using DIAMOND [24] against UniRef100 (state Dec. 2017) [25], modified with NCBI taxonomic information of the respective protein sequences (FunTaxDB, tentatively accessible through https://uni-duisburg-essen.sciebo.de/s/pi4cuYwyZ3KJVMl). The consensus taxonomy of each scaffold was predicted by considering the taxonomic rank of each protein on the scaffold on each taxonomic level and choosing the lowest taxonomic rank when more than 50% of the protein taxonomies agree. Reads were mapped to scaffolds using Bowtie2 [26] and the average scaffold coverage was estimated along with scaffolds’ length and GC content. Previously published ubiquitous single copy genes [27] were identified using HHmer 3.2 [28] and custom tables collecting GC, coverage, length, taxonomy and presence / absence of single copy genes of scaffolds were generated using scripts available along with uBin under https://github.com/ProbstLab/uBin-helperscripts.

#### Binning and curation

ISS assemblies were binned using Emergent Self-Organizing Maps (ESOM) [8]. Scaffolds were fragmented using the esomWrapper.pl [8] script, using 10kbp and 5kbp as maximum and minimum fragment sizes respectively. *Streptomyces griseus* NBRC13350 (high GC, NC_010572.1) and *Escherichia coli* K12 (low GC, NC_000913.3) genomes were spiked in to verify successful ESOM training. For ESOM training, the starting radius was set to 50 and the map-size was adjusted to the suggested size in the esomWrapper.pl output. ISS data was additionally binned directly using uBin.

CAMI datasets were binned using the automatic binners abawaca [29] and MaxBin2 [7], using both 3 kbp and 5 kbp as well as 5kbp and 10kbp as minimum and maximum fragment sizes respectively as abawaca input and using both available marker gene sets of MaxBin2 for binning. The output of the four different binners was aggregated using DAS Tool [6]. Tomsk and SulCav binning has been described previously [30].

Tables containing Bin, GC, coverage, length, taxonomy and single copy gene presence / absence information were loaded into uBin and used to curate draft genomes. Coding regions and single copy genes on genomes were predicted as described, omitting the -meta flag in prodigal.

#### Calculation of in situ replication indices

Bacterial *in situ* replication indices (iRep [1]) were calculated by mapping reads on the genomes and filtered for 3 mismatches, which correspond to 2% mismatch rate in the 150 bp reads. The rest of the settings for the iRep software were default.

#### Estimation of sample complexity

The sample complexity was estimated using the diversity of the *rpS3* marker gene. *rpS3* genes were annotated as described above. We are aware that sample complexity can also stem from other factors like K-mer frequency or coverage distribution patterns that this estimation does not take into account. However, these metrics cannot be assessed for environmental samples easily as the real composition is unknown. See *Tab. S4* for *rpS3* based complexity estimates across analyzed samples.

#### Phylogenomics

Ribosomal proteins were identified with blastp [31] (e-value 10^−5^) against 16 ribosomal proteins set as used in [32], aligned using muscle [33] with default parameters, trimmed with BMGE [34] and the BLOSUM30 substitution matrix and concatenated. The phylogenetic tree was calculated using Fasttree 2.1.8 [35] with default parameters. The tree was visualized in Dendroscope 3.7.2 [36].

#### Calculation of F-scores

Precision and recall of CAMI bins were determined using the known genomic assignment of the scaffolds and where they were allocated to during binning and curation. Genomic bins were assigned as corresponding to a CAMI genome based on the maximum scaffolds belonging to the same CAMI genome. Precision and recall of genomes from real-world datasets were determined using completeness as a proxy for True Positives, 1-%completeness as False Negatives, contamination as a proxy for False Positives and 1-contamination as True Negatives. The F-score was calculated as the mean between precision and recall.

#### Statistical evaluation

Statistical evaluation was performed in R [37]. Both paired and unpaired Welch t-tests [38] as well as Kruskal-Wallis [39] tests, one- and two-way ANOVA’s [40] and TukeyHSD [41] significance tests were performed. ggplot2 [42] was used to visualize data. The TOSTpaired.raw function within the TOSTER [43] package was used to confirm the non-significance of PMA-related tests, using 0.1 as the equivalence bound.

#### Metagenome availability

Accessions to raw reads and assemblies used in this study are listed in *Tab. S4*.

#### Software availability

The platform-independent genome curation software uBin is freely available under the MIT license at https://github.com/ProbstLab/uBin. The installation of the software from the OS-dedicated installers is dependency-free, while source code installation requires a Unix-based OS and package managers like npm or yarn.

**The authors declare no competing interest. All data is publicly available**

## Supplementary Figures

**Fig. S1.**
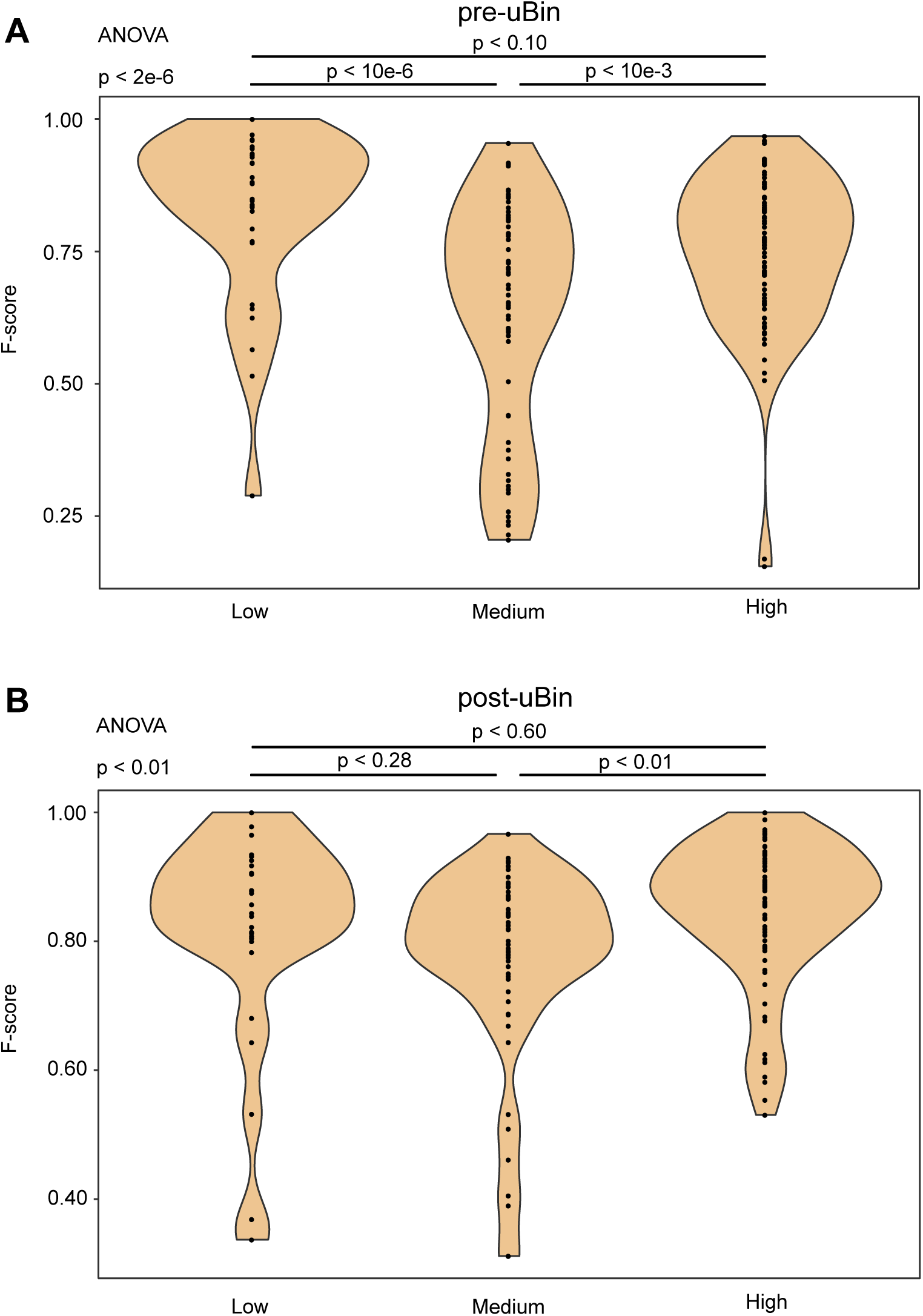
Comparison of bin qualities before and after uBin curation in simulated datasets with varying complexity. Compared are the bin qualities within bins before uBin (A) and within bins after uBin (B) in different complexities. ANOVA followed by the TukeyHSD post-hoc test were used to identify significant differences in quality between complexity groups.

**Fig. S2.**
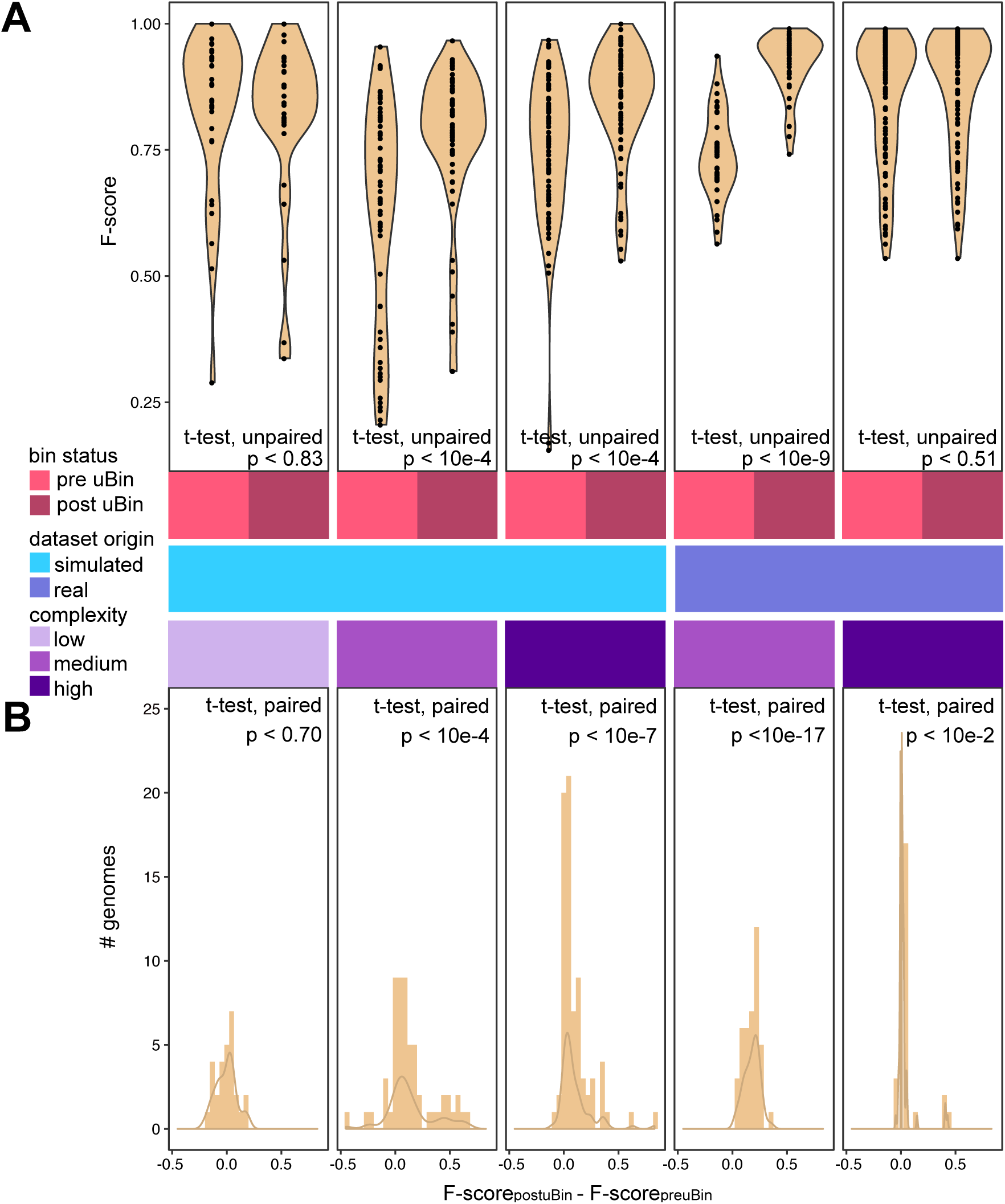
Performance of uBin on simulated and real datasets with varying degrees of complexity. **A:** Violin plots of the F-score (mean between recall and precision) of genomes prior to uBin curation (pre uBin) and after uBin curation (post uBin) across simulated low, medium and high complexity datasets of the CAMI challenge as well as real world metagenomic datasets of medium (Tomsk) and high (SulCav AS07-7) complexity. Unpaired Welch t-test p values are depicted. **B:** Histograms of the F-score differences for each bin prior and post uBin curation and their density distribution. Paired Welch t-test p-values are depicted.

**Fig. S3.**
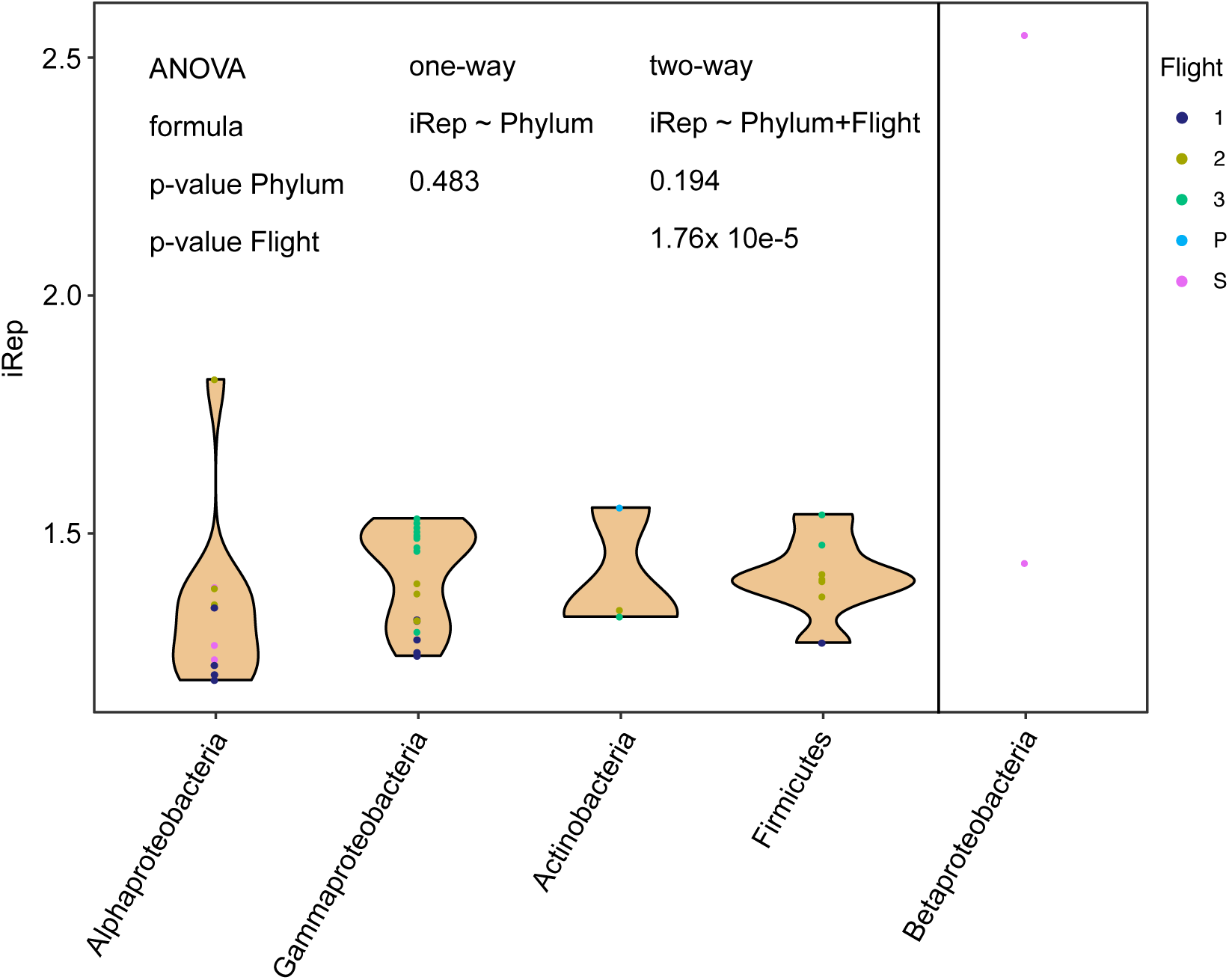
iRep distribution of ISS genomes by phylum. One- and two-way ANOVA were performed for significance testing. The Betaproteobacteria were excluded for the statistical analyses because of too few and highly diverging datapoints. No significant influence of the species was determined while the flight of origin was a significant coefficient in the two-way ANOVA analyses.

## Supplementary Tables

**Tab. S1 | F-scores pre- and post-uBin of CAMI, Tomsk and SulCav datasets.**

TableS1_Fscores_CAMI_Tomsk_SulCav.xlsx

**Tab. S2.**
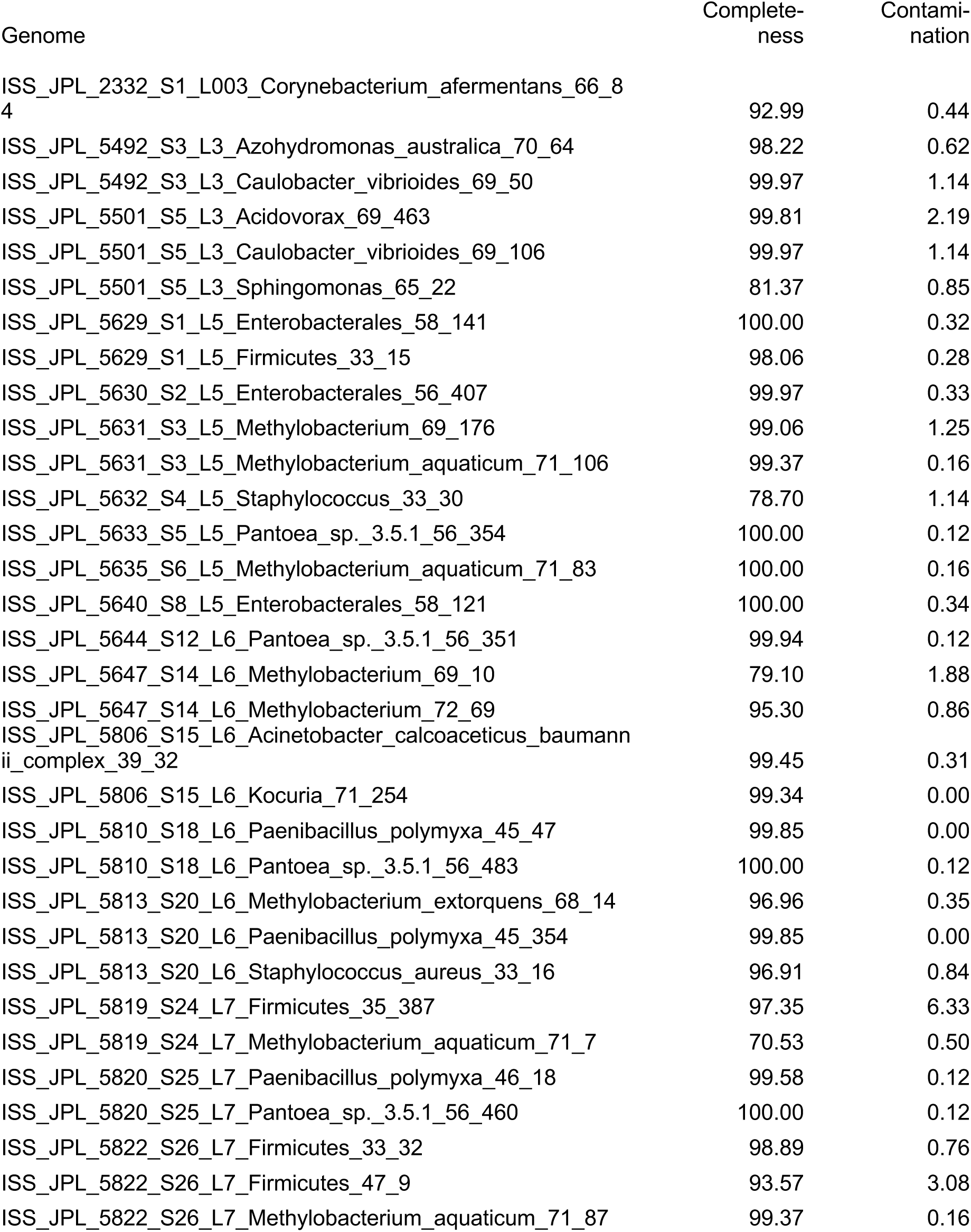
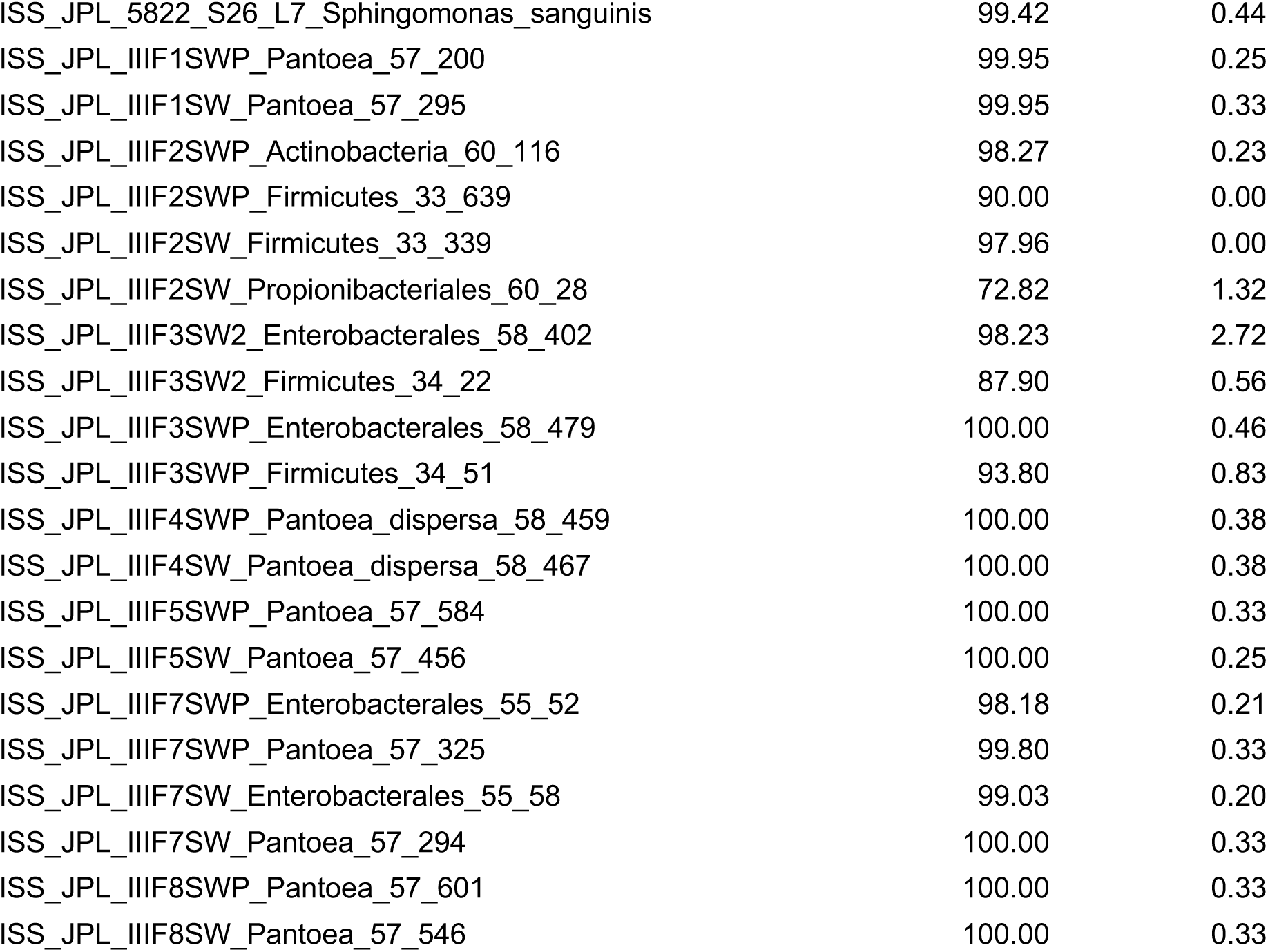
Genome statistics of recovered ISS genomes from metagenomes based on CheckM [14].

**Tab. S3.**
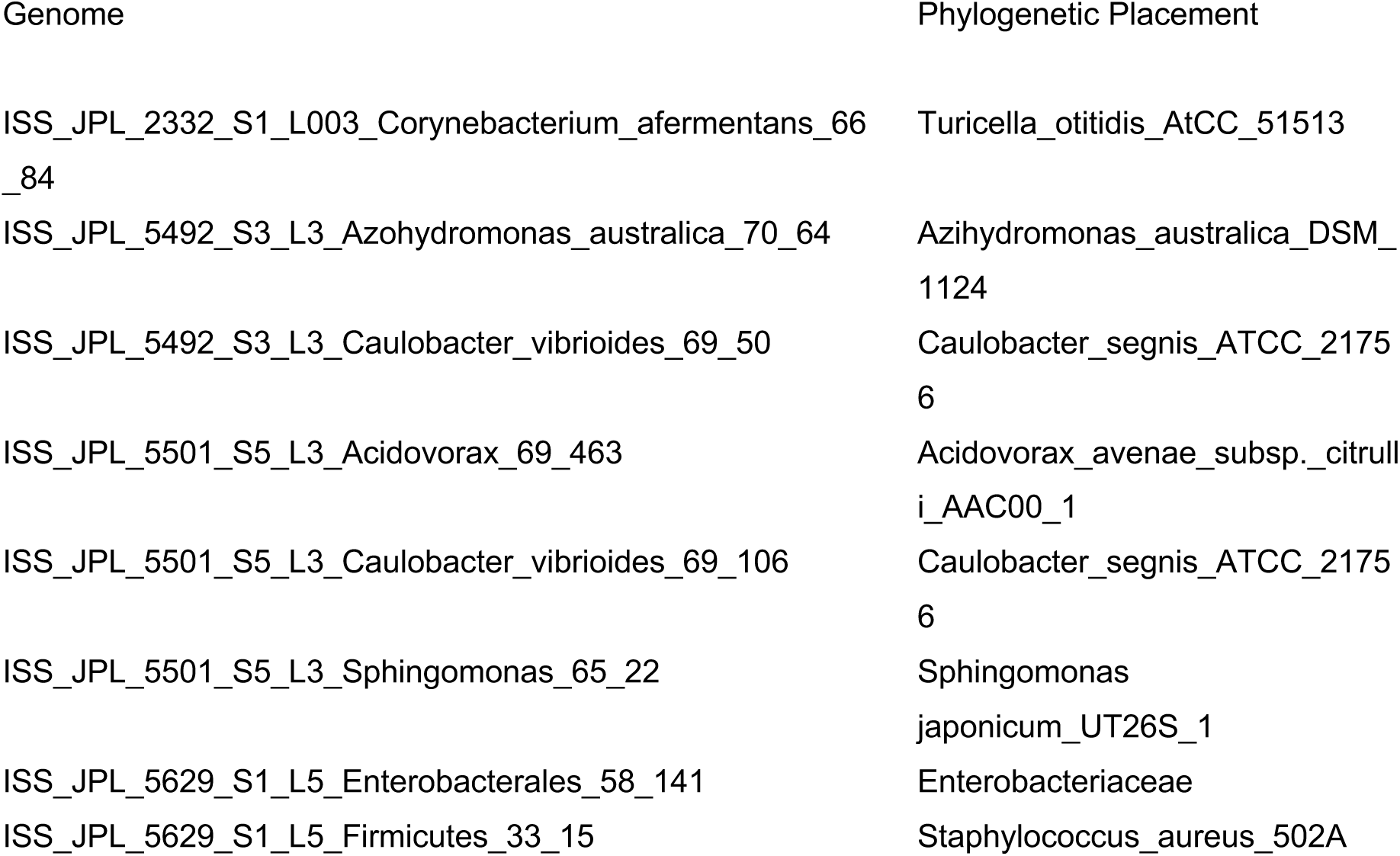
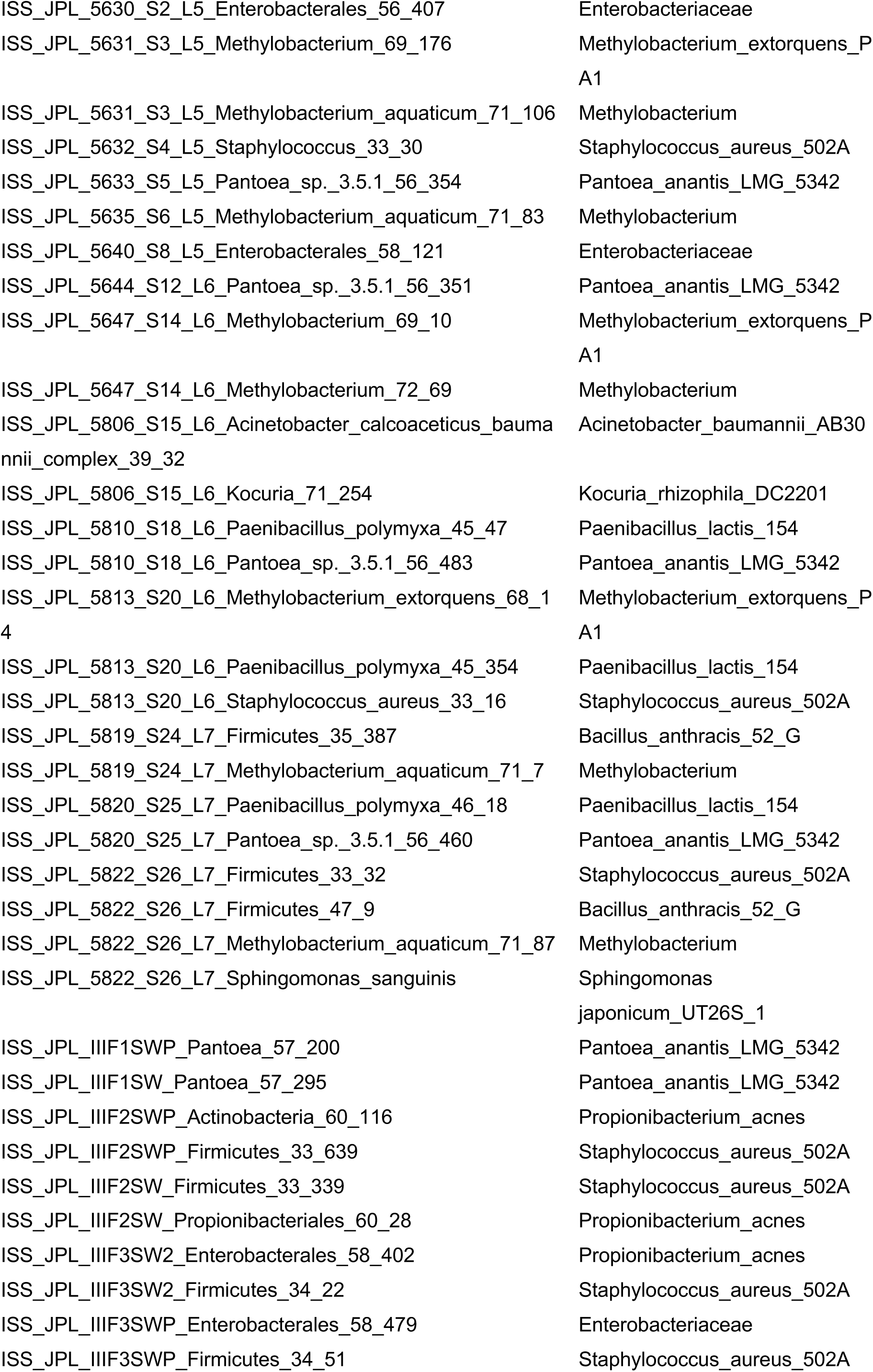
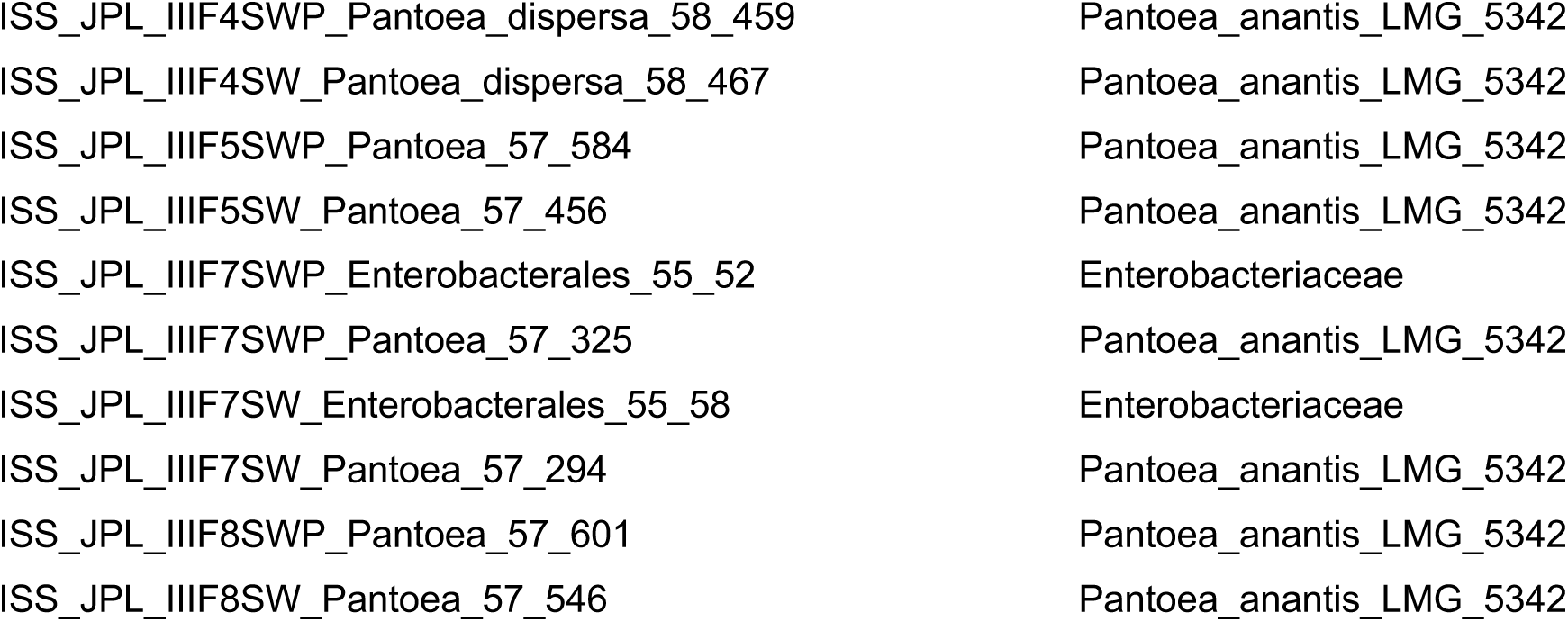
Phylogenetic characterization of recovered genomes. 16 ribosomal proteins were used to phylogenetically place the genomes. The taxonomy of the genomes represents the consensus taxonomy.

**Tab. S4.** Sample accessions and complexity based on the *rpS3* marker gene. Simulated low, medium and high complexity assembly and read datasets from the 1st CAMI challenge were downloaded from https://data.cami-challenge.org/participate. Raw reads from ISS flights I and II can be downloaded from the GeneLabs website (https://genelab-data.ndc.nasa.gov/genelab/accession/GLDS-66/).

| Sample | Read Accession | # rpS3 genes | Complexity category |
| --- | --- | --- | --- |
| CAMI_low | See table caption | 30 | Low |
| CAMI_medium_S1 | See table caption | 98 | Medium |
| CAMI_high_S1 | See table caption | 404 | High |
| Tomsk | SRR7102746 | 46 | Low-Medium |
| SulCav_AS07-7 | SRR1559028 | 151 | Medium-High |
| ISS_JPL_2332_S1_L003 | See table caption | 9 | Low |
| ISS_JPL_2333_S2_L003 | See table caption | 1 | Low |
| ISS_JPL_5492_S3_L003 | See table caption | 7 | Low |
| ISS_JPL_5501_S5_L003 | See table caption | 10 | Low |
| ISS_JPL_5629_S1_L005 | See table caption | 4 | Low |
| ISS_JPL_5630_S2_L005 | See table caption | 1 | Low |
| ISS_JPL_5631_S3_L005 | See table caption | 2 | Low |
| ISS_JPL_5632_S4_L005 | See table caption | 3 | Low |
| ISS_JPL_5633_S5_L005 | See table caption | 1 | Low |
| ISS_JPL_5635_S6_L005 | See table caption | 4 | Low |
| ISS_JPL_5640_S8_L005 | See table caption | 4 | Low |
| ISS_JPL_5644_S12_L006 | See table caption | 1 | Low |
| ISS_JPL_5647_S14_L006 | See table caption | 2 | Low |
| ISS_JPL_5806_S15_L006 | See table caption | 5 | Low |
| ISS_JPL_5808_S16_L006 | See table caption | 5 | Low |
| ISS_JPL_5810_S18_L006 | See table caption | 2 | Low |
| ISS_JPL_5813_S20_L006 | See table caption | 3 | Low |
| ISS_JPL_5818_S23_L007 | See table caption | 2 | Low |
| ISS_JPL_5819_S24_L007 | See table caption | 5 | Low |
| ISS_JPL_5820_S25_L007 | See table caption | 2 | Low |
| ISS_JPL_5822_S26_L007 | See table caption | 4 | Low |
| ISS_JPL_IIIF1SWP | SRX3808505 | 1 | Low |
| ISS_JPL_IIIF1SW | SRX3808512 | 1 | Low |
| ISS_JPL_IIIF2SWP | SRX3808504 | 3 | Low |
| ISS_JPL_IIIF2SW | SRX3808511 | 2 | Low |
| ISS_JPL_IIIF3SWP | SRX3808503 | 2 | Low |
| ISS_JPL_IIIF3SW | SRX3808514 | 2 | Low |
| ISS_JPL_IIIF4SWP | SRX3808508 | 1 | Low |
| ISS_JPL_IIIF4SW | SRX3808513 | 1 | Low |
| ISS_JPL_IIIF5SWP | SRX3808507 | 1 | Low |
| ISS_JPL_IIIF5SW | SRX3808510 | 1 | Low |
| ISS_JPL_IIIF7SWP | SRX3808529 | 1 | Low |
| ISS_JPL_IIIF7SW | SRX3808509 | 1 | Low |
| ISS_JPL_IIIF8SWP | SRX3808530 | 1 | Low |
| ISS_JPL_IIIF8SW | SRX3808535 | 1 | Low |

**Tab. S5.** ISS metagenome assembly statistics. Assembly statistics for SulCav AS07-7 and Tomsk metagenomes have been previously reported [30].

| Sample | #Gbp reads<br>after QC | #Mbp<br>scaffolds | #Mbp scaffolds ><br>1Kbp length | N50 scaffolds ><br>1Kbp length |
| --- | --- | --- | --- | --- |
| ISS_JPL_2332_S1_L003 | 9.5 | 859.2 | 54.7 | 1335 |
| ISS_JPL_2333_S2_L003 | 0.3 | 4.3 | 0.4 | 1344 |
| ISS_JPL_5492_S3_L003 | 6.7 | 87.6 | 65.9 | 10284 |
| ISS_JPL_5501_S5_L003 | 6.5 | 84.5 | 62.7 | 7814 |
| ISS_JPL_5629_S1_L005 | 3.6 | 127.2 | 76.4 | 38542 |
| ISS_JPL_5630_S2_L005 | 2.1 | 13.7 | 10.2 | 19268 |
| ISS_JPL_5631_S3_L005 | 3.6 | 68.5 | 22.5 | 51306 |
| ISS_JPL_5632_S4_L005 | 1.9 | 82.4 | 40.4 | 38085 |
| ISS_JPL_5633_S5_L005 | 2.5 | 57.6 | 30.1 | 1959 |
| ISS_JPL_5635_S6_L005 | 2.6 | 112.0 | 84.5 | 24153 |
| ISS_JPL_5640_S8_L005 | 2.7 | 88.5 | 58.4 | 29537 |
| ISS_JPL_5644_S12_L006 | 2.6 | 62.6 | 40.6 | 3170 |
| ISS_JPL_5647_S14_L006 | 3.5 | 77.5 | 64.9 | 9487 |
| ISS_JPL_5806_S15_L006 | 4.1 | 71.0 | 66.7 | 31164 |
| ISS_JPL_5808_S16_L006 | 3.4 | 128.2 | 55.8 | 53142 |
| ISS_JPL_5810_S18_L006 | 3.2 | 11.8 | 10.8 | 179991 |
| ISS_JPL_5813_S20_L006 | 2.6 | 53.6 | 16.1 | 17216 |
| ISS_JPL_5818_S23_L007 | 3.4 | 54.2 | 52.9 | 86659 |
| ISS_JPL_5819_S24_L007 | 2.6 | 64.8 | 40.0 | 3059 |
| ISS_JPL_5820_S25_L007 | 2.8 | 11.4 | 10.8 | 62007 |
| ISS_JPL_5822_S26_L007 | 2.1 | 29.0 | 18.9 | 41732 |
| ISS_JPL_IIIF1SWP | 1.0 | 7.3 | 5.4 | 132556 |
| ISS_JPL_IIIF1SW | 1.5 | 9.1 | 6.1 | 225749 |
| ISS_JPL_IIIF2SWP | 4.2 | 53.7 | 37.0 | 3007 |
| ISS_JPL_IIIF2SW | 2.1 | 36.0 | 10.2 | 1910 |
| ISS_JPL_IIIF3SWP | 2.7 | 14.4 | 7.5 | 168978 |
| ISS_JPL_IIIF3SW | 2.4 | 14.1 | 7.4 | 168987 |
| ISS_JPL_IIIF4SWP | 2.3 | 5.1 | 5.0 | 552627 |
| ISS_JPL_IIIF4SW | 2.4 | 5.8 | 5.1 | 405632 |
| ISS_JPL_IIIF5SWP | 2.9 | 5.1 | 4.9 | 205914 |
| ISS_JPL_IIIF5SW | 2.4 | 5.5 | 4.9 | 375929 |
| ISS_JPL_IIIF7SWP | 2.0 | 10.5 | 10.0 | 61535 |
| ISS_JPL_IIIF7SW | 1.8 | 11.4 | 10.0 | 59237 |
| ISS_JPL_IIIF8SWP | 3.0 | 6.2 | 5.0 | 248062 |
| ISS_JPL_IIIF8SW | 2.8 | 7.0 | 5.0 | 274906 |

## Additional Supplementary Files

**File S1 | Phylogenetic tree for placement of ISS genomes based on 16 ribosomal proteins.**

FileS1_ISS_PhyloTree.tree

## References

1. Brown CT, Olm MR, Thomas BC, Banfield JF. Measurement of bacterial replication rates in microbial communities. Nat Biotechnol. 2016;34:1256–63.

2. Moss EL, Maghini DG, Bhatt AS. Complete, closed bacterial genomes from microbiomes using nanopore sequencing. Nat Biotechnol. 2020;38:1–7.

3. Singleton CM, Petriglieri F, Kristensen JM, Kirkegaard RH, Michaelsen TY, Andersen MH, et al. Connecting structure to function with the recovery of over 1000 high-quality activated sludge metagenome-assembled genomes encoding full-length rRNA genes using long-read sequencing. bioRxiv. 2020;2020.05.12.088096.

4. Teeling H, Meyerdierks A, Bauer M, Amann R, Glöckner FO. Application of tetranucleotide frequencies for the assignment of genomic fragments. Environ Microbiol. 2004;6:938–47.

5. Albertsen M, Hugenholtz P, Skarshewski A, Nielsen KL, Tyson GW, Nielsen PH. Genome sequences of rare, uncultured bacteria obtained by differential coverage binning of multiple metagenomes. Nat Biotechnol. 2013;31:533–8.

6. Sieber CMK, Probst AJ, Sharrar A, Thomas BC, Hess M, Tringe SG, et al. Recovery of genomes from metagenomes via a dereplication, aggregation and scoring strategy. Nat Microbiol. 2018;3:836–43.

7. Wu Y-W, Simmons BA, Singer SW. MaxBin 2.0: an automated binning algorithm to recover genomes from multiple metagenomic datasets. Bioinformatics. 2016;32:605–7.

8. Dick GJ, Andersson AF, Baker BJ, Simmons SL, Thomas BC, Yelton AP, et al. Community-wide analysis of microbial genome sequence signatures. Genome Biol. 2009;10:R85.

9. Alneberg J, Bjarnason BS, de Bruijn I, Schirmer M, Quick J, Ijaz UZ, et al. Binning metagenomic contigs by coverage and composition. Nat Methods. 2014;11:1144–6.

10. Kang DD, Li F, Kirton E, Thomas A, Egan R, An H, et al. MetaBAT 2: an adaptive binning algorithm for robust and efficient genome reconstruction from metagenome assemblies. PeerJ. 2019;7:e7359.

11. Shaiber A, Eren AM. Composite Metagenome-Assembled Genomes Reduce the Quality of Public Genome Repositories. mBio. 2019;10.

12. Ballenghien M, Faivre N, Galtier N. Patterns of cross-contamination in a multispecies population genomic project: detection, quantification, impact, and solutions. BMC Biol. 2017;15:25.

13. Bowers RM, Kyrpides NC, Stepanauskas R, Harmon-Smith M, Doud D, Reddy TBK, et al. Minimum information about a single amplified genome (MISAG) and a metagenome-assembled genome (MIMAG) of bacteria and archaea. Nat Biotechnol. 2017;35:725–31.

14. Parks DH, Imelfort M, Skennerton CT, Hugenholtz P, Tyson GW. CheckM: assessing the quality of microbial genomes recovered from isolates, single cells, and metagenomes. Genome Res. 2015;25:1043–55.

15. Eren AM, Esen ÖC, Quince C, Vineis JH, Morrison HG, Sogin ML, et al. Anvi’o: an advanced analysis and visualization platform for ‘omics data. PeerJ. 2015;3:e1319.

16. Wrighton KC, Thomas BC, Sharon I, Miller CS, Castelle CJ, VerBerkmoes NC, et al. Fermentation, hydrogen, and sulfur metabolism in multiple uncultivated bacterial phyla. Science. 2012;337:1661–5.

17. Brinkrolf K, Schneider J, Knecht M, Rückert C, Tauch A. Draft genome sequence of Turicella otitidis ATCC 51513, isolated from middle ear fluid from a child with otitis media. J Bacteriol. 2012;194:5968–9.

18. Vallenet D, Engelen S, Mornico D, Cruveiller S, Fleury L, Lajus A, et al. MicroScope: a platform for microbial genome annotation and comparative genomics. Database (Oxford). 2009;2009.

19. Kanehisa M, Sato Y. KEGG Mapper for inferring cellular functions from protein sequences. Protein Sci. 2020;29:28–35.

20. Chen L-X, Anantharaman K, Shaiber A, Eren AM, Banfield JF. Accurate and complete genomes from metagenomes. Genome Res. 2020;30:315–33.

21. JN Fass NJ. Sickle: A sliding-window, adaptive, quality-based trimming tool for FastQ files. https://github.com/najoshi/sickle; 2011.

22. Nurk S, Meleshko D, Korobeynikov A, Pevzner PA. metaSPAdes: a new versatile metagenomic assembler. Genome Res. 2017;27:824–34.

23. Hyatt D, Chen G-L, LoCascio PF, Land ML, Larimer FW, Hauser LJ. Prodigal: prokaryotic gene recognition and translation initiation site identification. BMC Bioinformatics. 2010;11:119.

24. Buchfink B, Xie C, Huson DH. Fast and sensitive protein alignment using DIAMOND. Nat Methods. 2015;12:59–60.

25. Suzek BE, Huang H, McGarvey P, Mazumder R, Wu CH. UniRef: comprehensive and non-redundant UniProt reference clusters. Bioinformatics. 2007;23:1282–8.

26. Langmead B, Salzberg SL. Fast gapped-read alignment with Bowtie 2. Nat Methods. 2012;9:357–9.

27. Probst AJ, Castelle CJ, Singh A, Brown CT, Anantharaman K, Sharon I, et al. Genomic resolution of a cold subsurface aquifer community provides metabolic insights for novel microbes adapted to high CO2 concentrations. Environ Microbiol. 2017;19:459–74.

28. Eddy SR. Accelerated Profile HMM Searches. PLoS Comput Biol. 2011;7:e1002195.

29. Brown CT, Hug LA, Thomas BC, Sharon I, Castelle CJ, Singh A, et al. Unusual biology across a group comprising more than 15% of domain Bacteria. Nature. 2015;523:208.

30. Bornemann TLV, Adam PS, Turzynski V, Schreiber U, Figueroa-Gonzalez PA, Rahlff J, et al. Geological degassing enhances microbial metabolism in the continental subsurface. bioRxiv. 2020;2020.03.07.980714.

31. Altschul SF, Gish W, Miller W, Myers EW, Lipman DJ. Basic local alignment search tool. J Mol Biol. 1990;215:403–10.

32. Hug LA, Baker BJ, Anantharaman K, Brown CT, Probst AJ, Castelle CJ, et al. A new view of the tree of life. Nat Microbiol. 2016;1:16048.

33. Edgar RC. MUSCLE: multiple sequence alignment with high accuracy and high throughput. Nucleic acids res. 2004;32:1792–1797.

34. Criscuolo A, Gribaldo S. BMGE (Block Mapping and Gathering with Entropy): a new software for selection of phylogenetic informative regions from multiple sequence alignments. BMC Evol Biol. 2010;10:210.

35. Price MN, Dehal PS, Arkin AP. FastTree: Computing Large Minimum Evolution Trees with Profiles instead of a Distance Matrix. Mol Biol Evol. 2009;26:1641–50.

36. Huson DH, Richter DC, Rausch C, Dezulian T, Franz M, Rupp R. Dendroscope: An interactive viewer for large phylogenetic trees. BMC Bioinformatics. 2007;8:460.

37. R Core Team. R: A Language and Environment for Statistical Computing. 2008;R Foundation for Statistical Computing, Vienna, Austria.

38. Welch BL. The generalisation of student’s problems when several different population variances are involved. Biometrika. 1947;34:28–35.

39. Kruskal WH, Wallis WA. Use of Ranks in One-Criterion Variance Analysis. J AM STAT ASSOC. 1952;47:583–621.

40. St»hle L, Wold S. Analysis of variance (ANOVA). Chemometrics Intell Lab Sys. 1989;6:259–72.

41. Haynes W. Tukey’s Test. In: Dubitzky W, Wolkenhauer O, Cho K-H, Yokota H, editors. Encyclopedia of Systems Biology. New York, NY: Springer New York; 2013. p. 2303–4.

42. Wickham H. ggplot2: Elegant Graphics for Data Analysis. New York: Springer-Verlag; 2009.

43. Lakens D, Scheel AM, Isager PM. Equivalence Testing for Psychological Research: A Tutorial: AMPPS. 2018;1:259–69.

